# Zebrafish *duox* mutations provide a model for human congenital hypothyroidism

**DOI:** 10.1101/372003

**Authors:** Kunal Chopra, Shoko Ishibashi, Enrique Amaya

## Abstract

Thyroid dyshormonogenesis is a leading cause of congenital hypothyroidism, a highly prevalent but treatable condition. Thyroid hormone synthesis is dependent on the formation of reactive oxygen species (ROS). In humans, the primary sources for ROS production during thyroid hormone synthesis are the NADPH oxidase, DUOX1 and DUOX2. Indeed mutations in *DUOX1* and *DUOX2* have been linked with congenital hypothyroidism. Unlike humans, zebrafish has a single orthologue for *DUOX1* and *DUOX2*. In this study, we investigated the phenotypes associated with two nonsense mutant alleles of the single *duox* gene in zebrafish, *sa9892* and *sa13017*. Both alleles gave rise to readily observable phenotypes reminiscent of congenital hypothyroidism, from the larval stages through to adulthood. By using various methods to examine the external and internal phenotypes, we discovered a strong correlation between TH synthesis and *duox* function, beginning from the early larval stage, when T_4_ levels are already noticeably absent in the mutants. Loss of T_4_ production resulted in growth retardation, pigmentation defects, ragged fins, thyroid hyperplasia / external goiter, and infertility. Remarkably all of these defects associated with chronic congenital hypothyroidism could be rescued with T_4_ treatment, even when initiated when the fish had already reached adulthood. Our work suggests that these zebrafish *duox* mutants may provide a powerful model to understand the aetiology of untreated and treated congenital hypothyroidism even in advance stages of development.

## INTRODUCTION

Congenital hypothyroidism (CH) is an endocrine disorder that may result from disrupted thyroid hormone (TH) synthesis (15-20% of all cases) or impaired development of the thyroid gland [thyroid dysgenesis (TD)] (80% of all cases) (Kizys et al., 2017). CH is the most prevalent congenital endocrine disorder and also believed to be one of the most preventable causes of mental retardation (Chakera et al., 2012; Olivieri, 2015; Roberts and Ladenson, 2004). Indeed, in infants younger than 3 months of age neurological damage progressively worsens with delay in starting treatment (Virtanen et al., 1983). Mutations in the NADPH oxidases, *DUOX2* and to a lesser extent *DUOX1* have been associated with dyshormonogenesis in CH patients (Aycan et al., 2017; Moreno et al., 2002). DUOX1 and DUOX2 generate hydrogen peroxide (H_2_O_2_), which is a crucial electron acceptor during thyroid peroxidase-catalyzed iodination and coupling reactions occurring while TH synthesis is underway (De Deken et al., 2000; Dupuy et al., 1999). H_2_O_2_ production is a limiting step in TH biosynthesis. The main source of H_2_O_2_ in the thyroid is DUOX2 in conjunction with its maturation factor DUOX2A, both of which are located at the apical surface of the thyroid follicular cells, thyrocytes. DUOX2-mediated H_2_O_2_ acts as a thyroperoxidase (TPO) co-substrate, rapidly oxidising iodine and resulting in the covalent binding of iodide to the tyrosine residues of thyroglobulin in the follicular lumen. This produces monoiodotyrosine (MIT) and diiodotyrosine (DIT), in the thyroglobulin molecule, which undergo coupling to give the THs triiodothyronine (T_3_) and thyroxine (T_4_) (Carvalho and Dupuy, 2013; Muzza and Fugazzola, 2017; Sugawara, 2014). A negative feedback loop is in charge of thyroid size and function. Thyrocytes secrete T_3_ and T_4;_ and T_3_ and T_4_ inhibit the production of the thyroid-stimulating hormone (TSH) via the anterior pituitary thyrotropes (Dumont et al., 1992). Thyrocytes respond to limiting physiological stimuli by way of hypertrophy and proliferation. This is a direct response to compensate for diminishing THs in conditions including, but not limited to, iodine deficiency, exposure to anti-thyroid drugs and punctuated production of reactive oxygen species (ROS). It has been shown that early initiation of TH treatment (within three weeks post-partum) leads to normal IQ and physical growth and correlates with excellent prognoses (Aronson et al., 1990; Clause, 2013; Rahmani et al., 2016; Rovet et al., 1987). Expectedly then, if treatment is delayed beyond 4 weeks, individuals become increasingly prone to mental retardation and incomplete physical growth (Gilbert et al., 2012; Zimmermann, 2011). To date, various approaches have been adopted to induce hypothyroidism in animal models, including surgical removal of the thyroid gland, thyroid gland removal via radioactive iodine isotope (^131^I), dietary restriction of iodine, and goitrogen administration (Argumedo et al., 2012). We present here a zebrafish model of CH, which exhibits several phenotypes associated with CH in humans, including growth retardation. Interestingly, while CH zebrafish display growth retardation initially, they are able to reach normal size eventually without the need for pharmacological intervention. The additional external and internal phenotypes associated with hypothyrodism are restored upon treatment with T_4_, including restoration of reproductive function, even when treatment is not initiated until adulthood.

## RESULTS

### Molecular Characterisation of *duox* Mutant Alleles

Duox is a member of the NADPH oxidase (NOX) family of enzymes. Seven NOX family members are present in the human genome: NOX1, NOX2, NOX3, NOX4, NOX5, DUOX1 and DUOX2, and their primary function is to produce reactive oxygen species (ROS). All NOX enzymes are transmembrane proteins, exhibiting structural and functional conservation. They participate in electron transport across biological membranes, effecting the reduction of molecular oxygen to superoxide (Bedard and Krause, 2007). All NOX enzymes share conserved structural domains, including intracellular C-terminal tails containing NADPH and FAD binding sites and six transmembrane domains anchoring four highly conserved heme-binding histidines. DUOXes have an additional transmembrane domain, an extracellular N-terminal domain with peroxidase homology and two EF Ca^2+^ binding hands within their first intracellular loop (Fig. 1A)(Rada and Leto, 2008). The zebrafish genome encodes a single *duox* gene, rather than two *DUOX* paralogues present in humans (*DUOX1* and *DUOX2*) and lacks a *NOX3* orthologue (Kawahara et al., 2007). In zebrafish *duox* is located on chromosome 25 and encodes a 1528 amino acid protein. In order to investigate the function of *duox* in zebrafish, we obtained two nonsense mutation alleles, which arose from a large-scale ENU mutagenesis screen (Kettleborough et al., 2013). One allele, *duox sa9892,* contains a nonsense mutation in exon 21, resulting in a C>T change (Fig. 1B) and a premature stop codon (TAG) after the 944th amino acid; the second allele, *duox sa13017*, contains a nonsense mutation in exon 23, resulting in a C>T change (Fig. 1C) creating a premature stop codon (TGA) after the 997^th^ amino acid. Since both these premature codons result in truncations of the Duox protein, including the loss of the two critical C-terminal NADPH and FAD binding sites, they would be expected to be loss-of-function mutations. Genotyping of these alleles can be performed via genomic PCR followed by sequencing of these regions (see Fig. 1B and 1C).

**Fig. 1.**
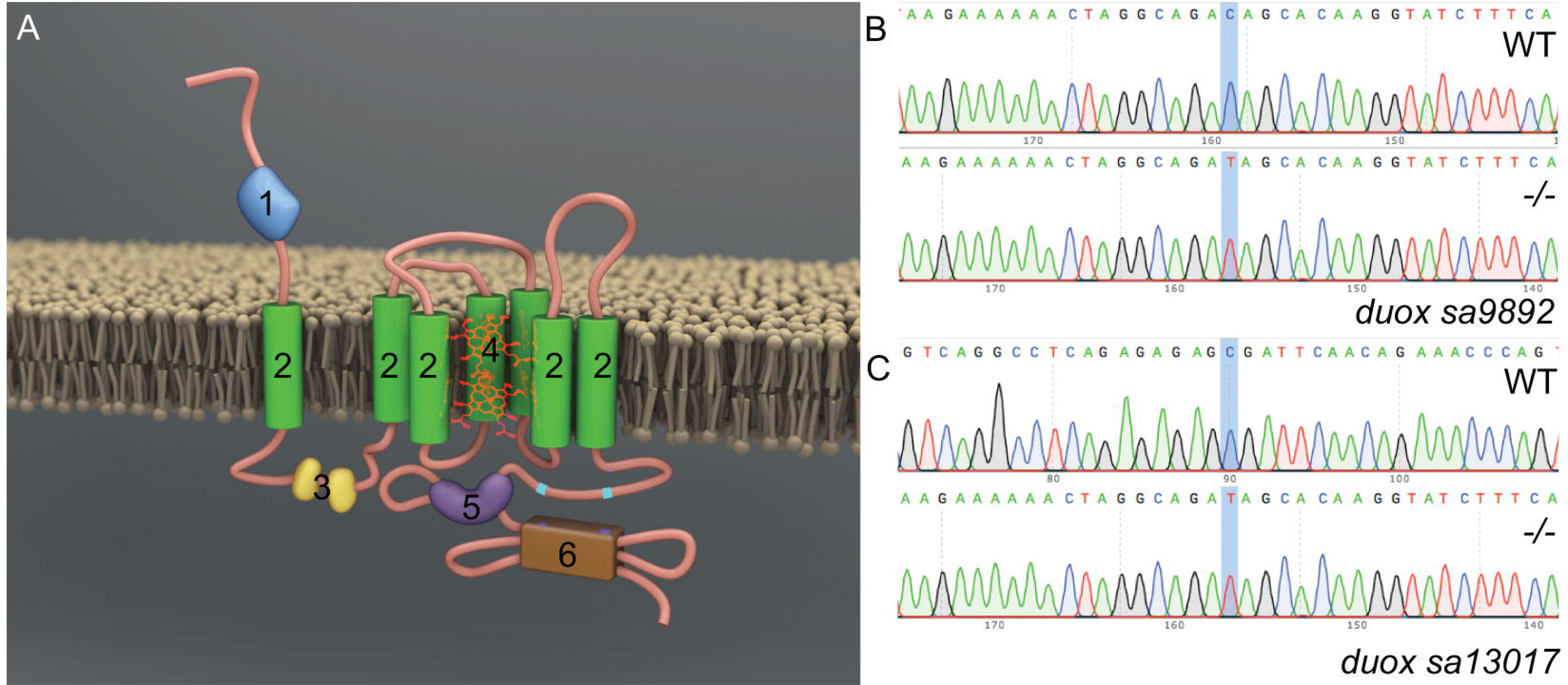
Molecular characterisation of *duox* mutant alleles. Duox is a transmembrane protein belonging to the NADPH oxidase family of enzymes. Duox has an NADPH oxidase domain at the C-terminus as well as a N-terminus peroxidase domain, thus named Dual oxidase (A). Characterisation of *duox sa9892* (B) and *duox sa13017* (C), via Sanger sequencing, shows the single nucleotide change C>T in contrast to a wild type reference sequence.

### *duox* Mutants are Growth Retarded

We in-crossed *duox sa9892^+/-^* and *duox sa13017^+/-^* sibling adults and inter-crossed *duox sa9892^+/-^* with *duox sa13017^+/-^* adults to produce a range of wild-type, heterozygous, homozygous mutant and compound heterozygous mutant animals containing both alleles. While the wild type, *duox sa9892^+/-^* and *duox sa13017^+/-^* animals were phenotypically indistinguishable, the homozygous mutants of both alleles, and the compound mutants (i.e. *duox sa9892/duox sa13017)* displayed a number of phenotypes that were distinct from those seen in the wild type and heterozygous siblings. The first overtly apparent phenotype exhibited by the *duox sa9892^-/-^*, the *duox sa13017^-/-^* and the trans-heterozygous *duox sa9892/duox sa13017* mutants was that they were growth retarded. At three months of age, the *sa9892^-/-^, sa13017^-/-^* and *sa9892/sa13017* mutant fish were significantly shorter in terms of body length than their WT and heterozygous siblings (Fig. 2A-2G). At six-months of age the *duox sa9892^-/-^* and *sa9892/sa13017* mutant animals caught up in size with their WT and heterozygous siblings. However, the *duox sa13017^-/-^* still remained stunted (Fig. 2H). Another phenotype suggestive of slowed growth was apparent in the growth and organogenesis of the swim bladder. The swim bladder is a hydrostatic organ, which becomes bi-lobed by 21 dpf (Winata et al., 2009). We found that the swim bladder in the homozygous *duox sa9892* mutant animals remained unilobed even at 54 dpf (Fig. 2L-2M). Homozygous *duox* mutants also exhibit a delay or absence of development of barbels, which are a set of anterior sensory appendages. Zebrafish develop a short pair of nasal barbels and a long trailing pair of maxillary barbels. These are normally visible by one month post-fertilisation and sustained throughout life (LeClair and Topczewski, 2010). In all cases, homozygous *duox* mutants lacked barbels at three months of age (Fig. 2I-2J). However, between 6 and 10 months of age, maxillary barbels were seen in some, but not all, older *duox sa9892^-/-^* (5 out of 11; see Fig. 2K) and *sa9892/sa13017* (2 out of 11) mutant animals, but in none of the *sa13017^-/-^* mutant animals (0 out of 6).

**Fig. 2.**
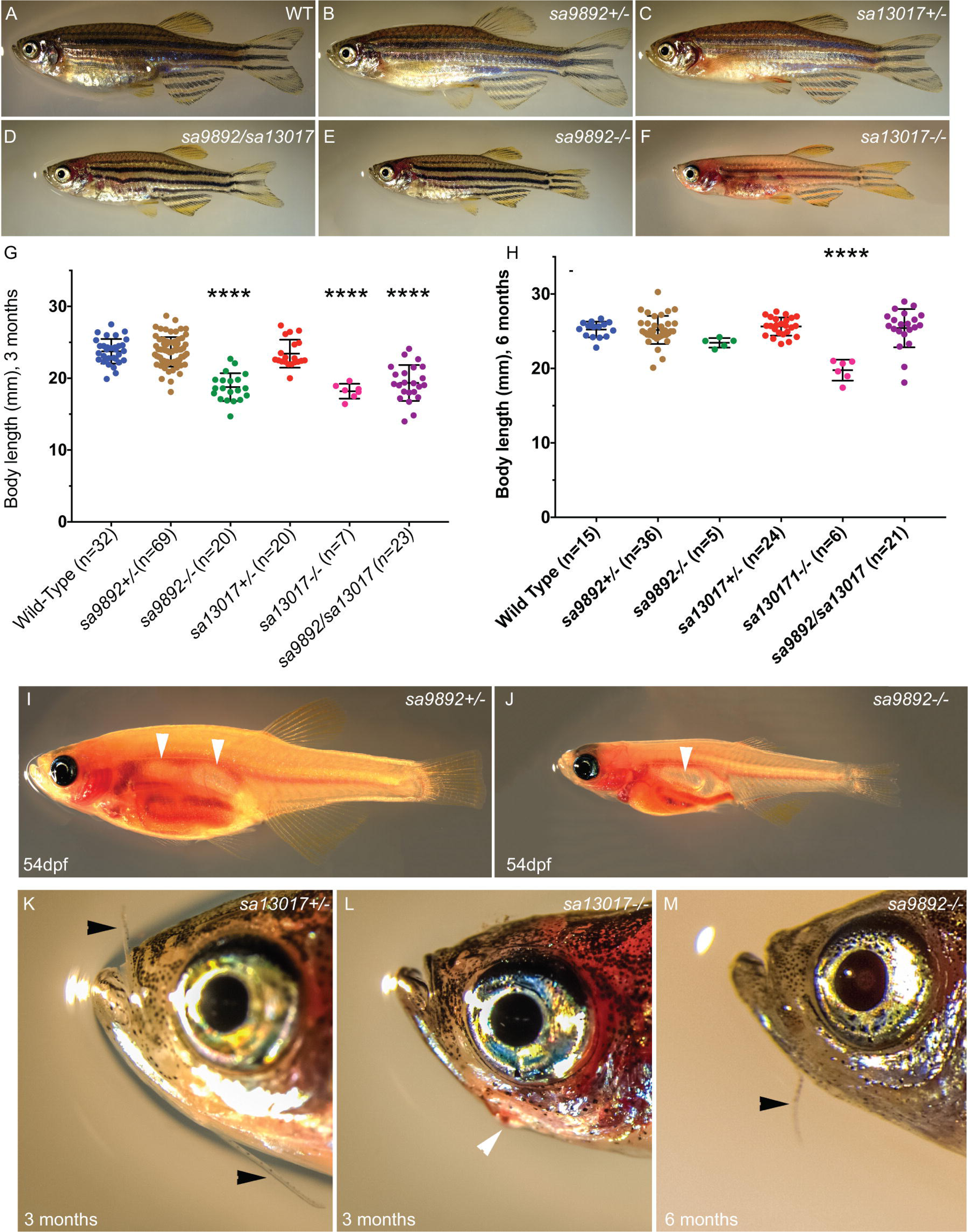
*duox* mutants exhibit growth retardation. Mutants for both alleles as well as compound heterozygotes are shorter than their wild type and heterozygous siblings at 3 months (A-G) but catch up by 6 months (H). *sa13017^-/-^* animals are trailing behind even at 6 months (H). Asterisks in G denote statistically significant differences (Bonferroni’s multiple comparisons test, **** P<0.0001) *duox* mutants also have a delay in the inflation of the anterior lobe of the swim bladder (I-J) (white arrows indicate lobes). Adults up to 6 months old also lack of barbels (dark arrows, K-M). Barbels emerge in some older animals (10months and above) (M). External goitres are often visible in young adults (white arrow, L).

### *duox* Mutants Have Dark Pigmentation, Erythema And Ragged Fins

Zebrafish are recognisable by their eponymous pattern of five dark blue stripes with alternating four lighter yellow inter-stripes, covering the lateral flanks, and anal and caudal fins (Singh and Nüsslein-Volhard, 2015). The dark blue stripes are comprised of black melanophores with a few iridescent iridophores, while yellow inter-stripes are comprised of yellow and orange xanthophores and numerous iridophores (Hirata et al., 2003). We found that the homozygous *duox* mutants were darker than their WT and heterozygote siblings (Fig. 3A-3C). The darker pigmentation was associated with the presence of approximately twice the number of melanophores in the homozygous *duox* mutants (*sa9892^-/-^ and sa9892/sa13017*) relative to their heterozygous and WT siblings (Fig. 3F). We found that the difference in pigmentation was sufficient to allow the visual identification of homozygous *duox* mutants from their heterozygous and wild type siblings, with 100% accuracy, as confirmed via genotyping. In addition, homozygous *duox* mutants also showed stripe irregularities not seen in WT and heterozygous siblings, such as wavy stripes and stripe discontinuities (Fig. 3D-3E). Thus pigmentation differences can be used as a reliable identification method for distinguishing homozygous *duox* mutants from their heterozygous and WT siblings as early as 60 dpf.

**Fig. 3.**
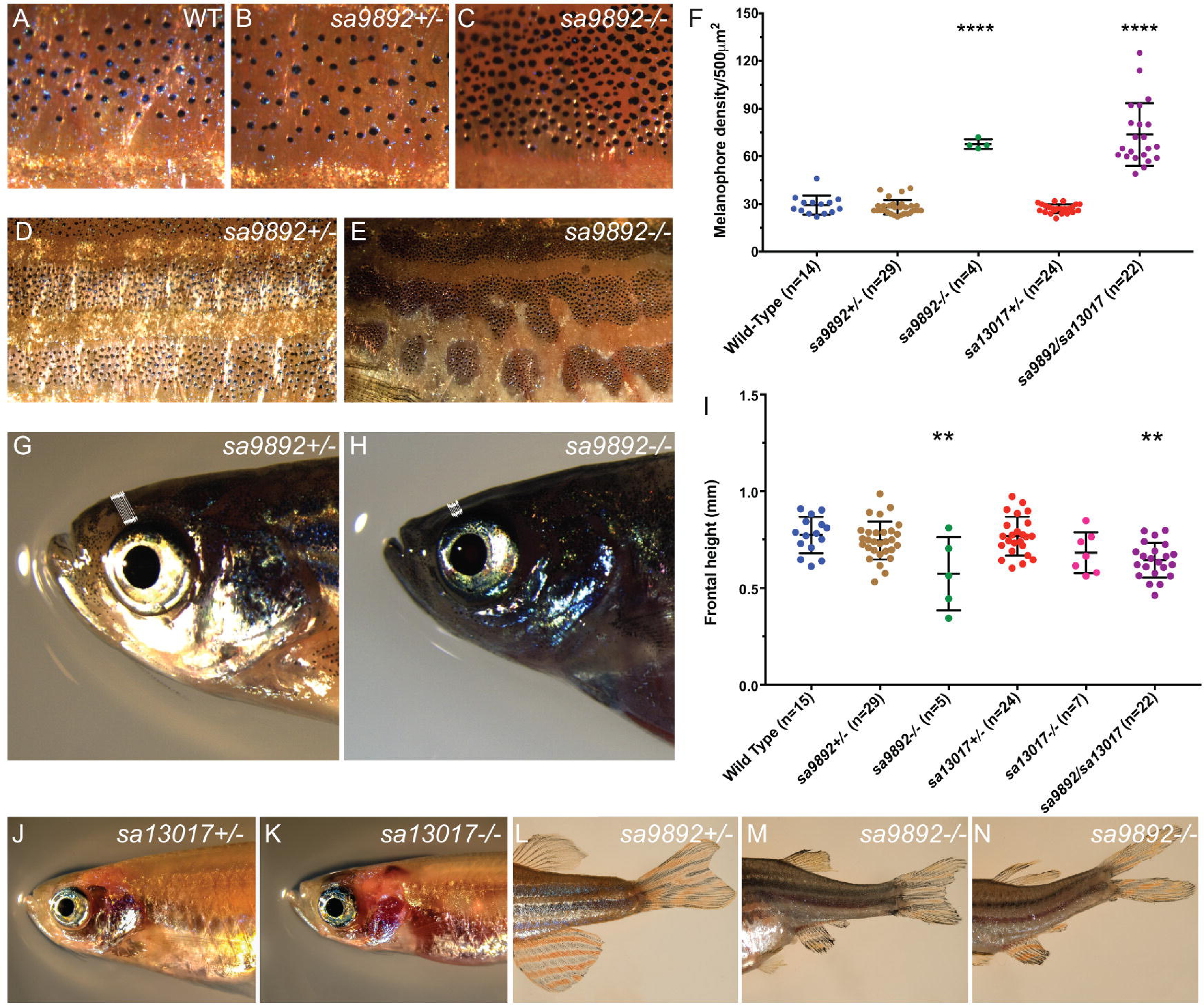
Adult *duox* mutant zebrafish display an array of visible phenotypes. A-C, 5X magnification of flank region showing the distribution of melanophores in wild type, *sa9892^+/-^* and *sa989^-/-^* siblings. The apparent abundance of melanophores was statistically significant in *duox* mutants (F). Asterisks denote statistically significant differences (Bonferroni’s multiple comparisons test, **** P<0.0001). *duox* mutants also showed irregularities in stripe pattern in contrast to heterozygous siblings, shown here in a 2X magnification of the flank in *sa9892* siblings (D-F). Craniofacial anomalies were evident among mutants, with frontal height significantly shorter among mutants (G–I) (Bonferroni’s multiple comparisons test, *P<0.5, ** P<0.01). Erythema in the thoracic region was prominent among mutants. This was especially noticeable in *nacre* backgrounds (J-K). *duox* mutants also suffered from perpetual fin damage, which manifest as ragged margins and tears (L-M).

Adding further to the list of phenotypes, we noticed erythema (redness) in the opercular region (Fig. 3J-3K). This was especially prominent in background strains that lack melanophores, such as *nacre*. The redness was most apparent in juvenile fish. Less apparent but nevertheless significant were craniofacial anomalies among adult mutants. In particular, we found a significant shortening of the frontal height among the *duox sa9892^-/-^* and *sa9892/sa13017* animals, when compared to their wild type and heterozygous siblings (Fig. 3G-3I).

Finally, the homozygous *duox* mutants often displayed misshapen or damaged fins. For example, we found that the *duox sa9892^-/-^* (15 out of 15), *sa13017^-/-^* (7 out of 7) and *sa9892/sa13017* (19 out of 23) animals displayed damaged fins. In many cases this was manifested as vertical (dorsal and anal fin) or horizontal (caudal fin) tears in the fins. In other cases, there were spontaneous losses of portions of fins or ragged fin margins (Fig. 3L-3N). Damaged fins were noticeable as early as 42 dpf.

### Homozyogous *duox* Mutants Are Viable But Are Unable To Breed

While we found that homozygous *duox* mutants reached adulthood, unlike their heterozygous and WT siblings, they were unable to breed. Females, although gravid, were found not to lay eggs regardless of pairing with mutant, heterozygous or WT males. Similarly, mutant males failed to cross with females, regardless of genotype. Furthermore, we noticed that homozygous *duox* mutant females seemed egg-bound, suggesting that they were unable lay eggs (Fig 4A-4B; left panels). We confirmed that females do contain eggs internally via histological sectioning (Fig 4A-4B; right panels) as well as via abdominal squeezing to release oocytes. Similarly, compound heterozygotes of the two alleles were found to be viable but failed to breed.

**Fig. 4.**
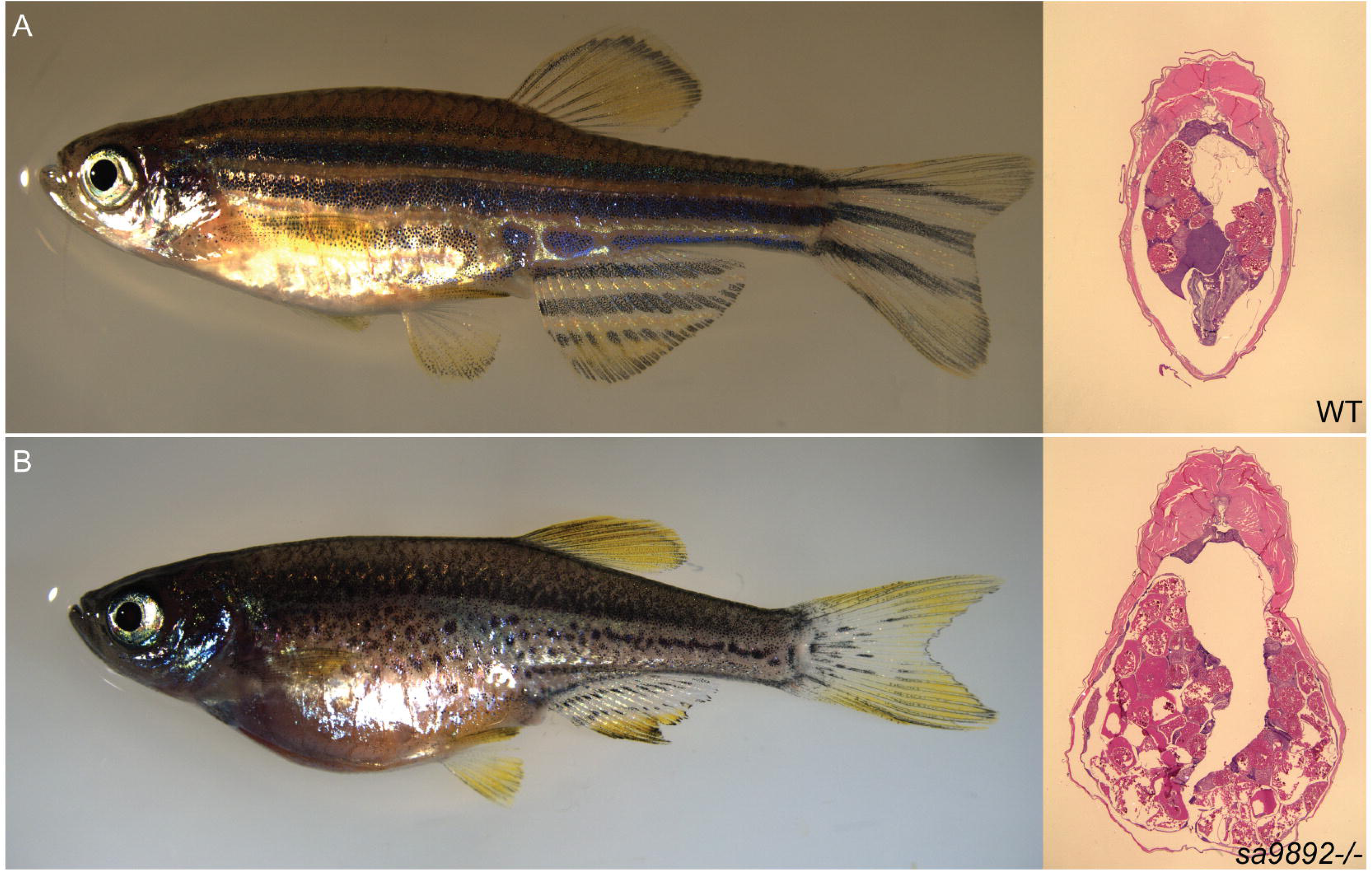
*duox* mutant females are unable to ovulate and become egg bound. H&E staining of abdominal sections reveals oocytes (A-B).

### Homozyogous *duox* Mutants Develop Goitres

In addition to the phenotypes described above, we noted that some of the homozygous *duox* mutant adult animals displayed goitre-like growths in the submandibular area (Fig. 5A-5B). These goitre-like growths were observed among all three mutant strains; *duox sa9892^-/-^* adults (11 out of 18), *sa13017^-/-^* adults (6 out of 11) and *sa9892/sa13017* adults (3 out of 22), which were older than 3 months of age. These richly vascularised growths were variously sized. Additionally, some of the animals also exhibited lateral flaring of opercular flaps (Fig. 5C). To confirm whether these goitre-like growths were indeed enlarged thyroids, we fixed and sectioned a subset of homozygous *duox* mutants and some of their wild type and heterozygous mutant siblings and performed ISH analysis for the expression of *thyroglobulin*, a thyroid marker. The results confirmed that these growths were indeed of thyroid origin (Fig. 5D, and 5G-5I). Also, the extent of thyroid hyperplasia was in striking contrast to the size of the thyroids in the WT and *sa9892^+/-^* siblings, where the extent of *thyroglobulin* staining was much smaller and more distinct, as discreet rings confined to the ventral mid-pharyngeal region (Fig. 5E-5F).

**Fig. 5.**
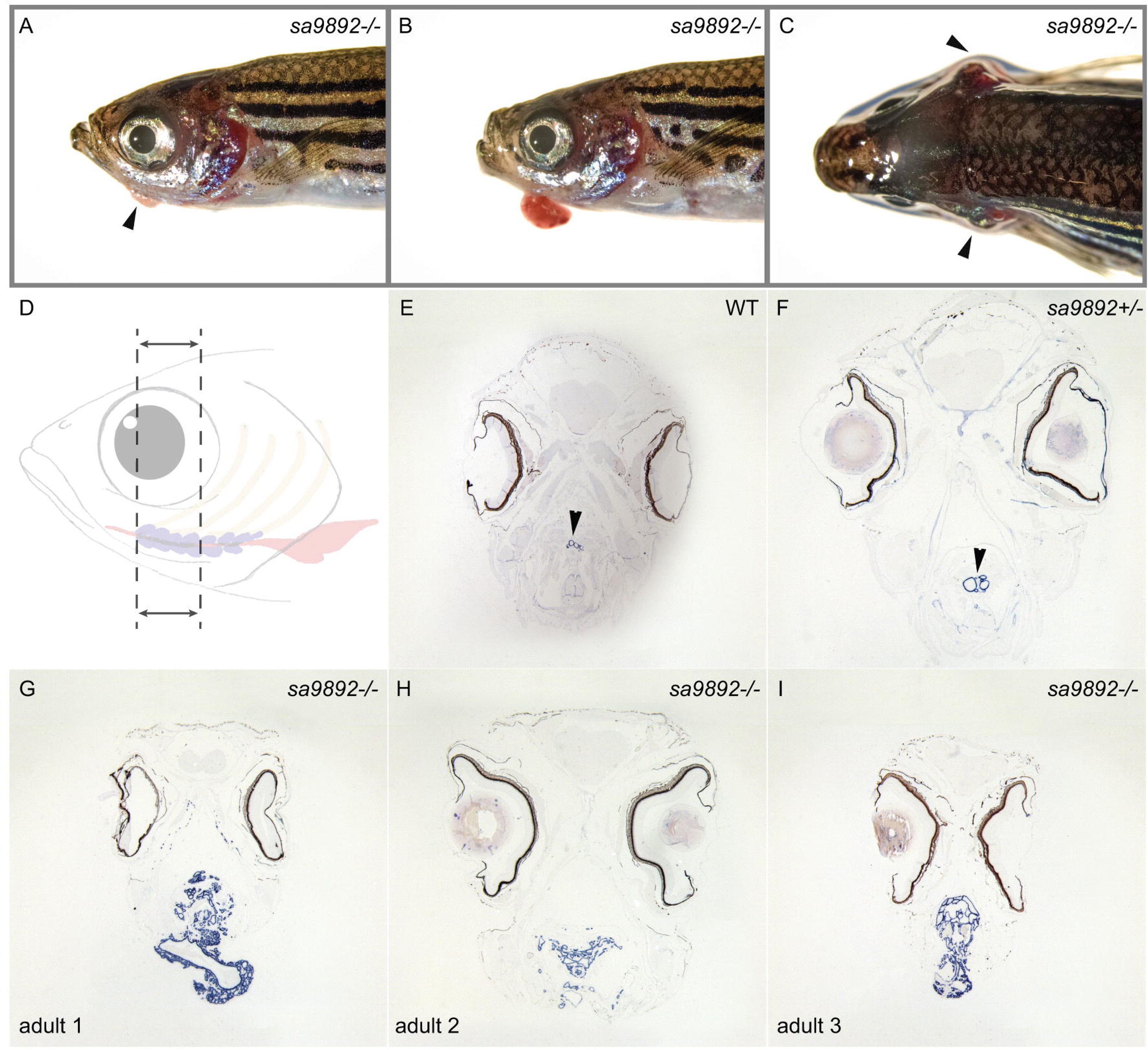
Homozygous *duox* mutations lead to goitre. Adult mutant animals exhibit an array of variably sized external goitres, as well as lateral flaring of opercula (A–C). When sectioned along the length of the follicular region (dotted area, D) and subjected to ISH for *thyroglobulin*, mutants reveal extensive spread of and ectopic thyroid follicular tissue (G–I), in contrast to the localised, discreet distribution in wild type and heterozygous siblings (E–F).

### *duox* Mutants Are In A State Of Hypothyroidism

The goitre-like growths, as well as the other phenotypes observed in the homozygous *duox* mutants suggested that the mutants might be exhibiting hypothyroidism. To test whether this might be the case, we assessed the presence of thyroxine (T_4_) in the homozygous *duox* mutants and in their heterozygous and WT siblings via whole-mount immunostaining. We found that, while the WT and heterozygous siblings exhibited robust T_4_ staining (Fig. 6A-6C), *duox sa9892^-/-^* and *sa13017^-/-^* larvae had no detectable T_4_ staining in their thyroids (Fig. 6D-6E). Consistent with the loss of T_4_ being due to lack of NADHP oxidase activity in the homozygous *duox* mutants, we were able to phenocopy the loss of T_4_ staining in the larvae by treating them with the NADPH oxidase inhibitor, diphenyleneiodonium (DPI) (Fig. 6F).

**Fig. 6.**
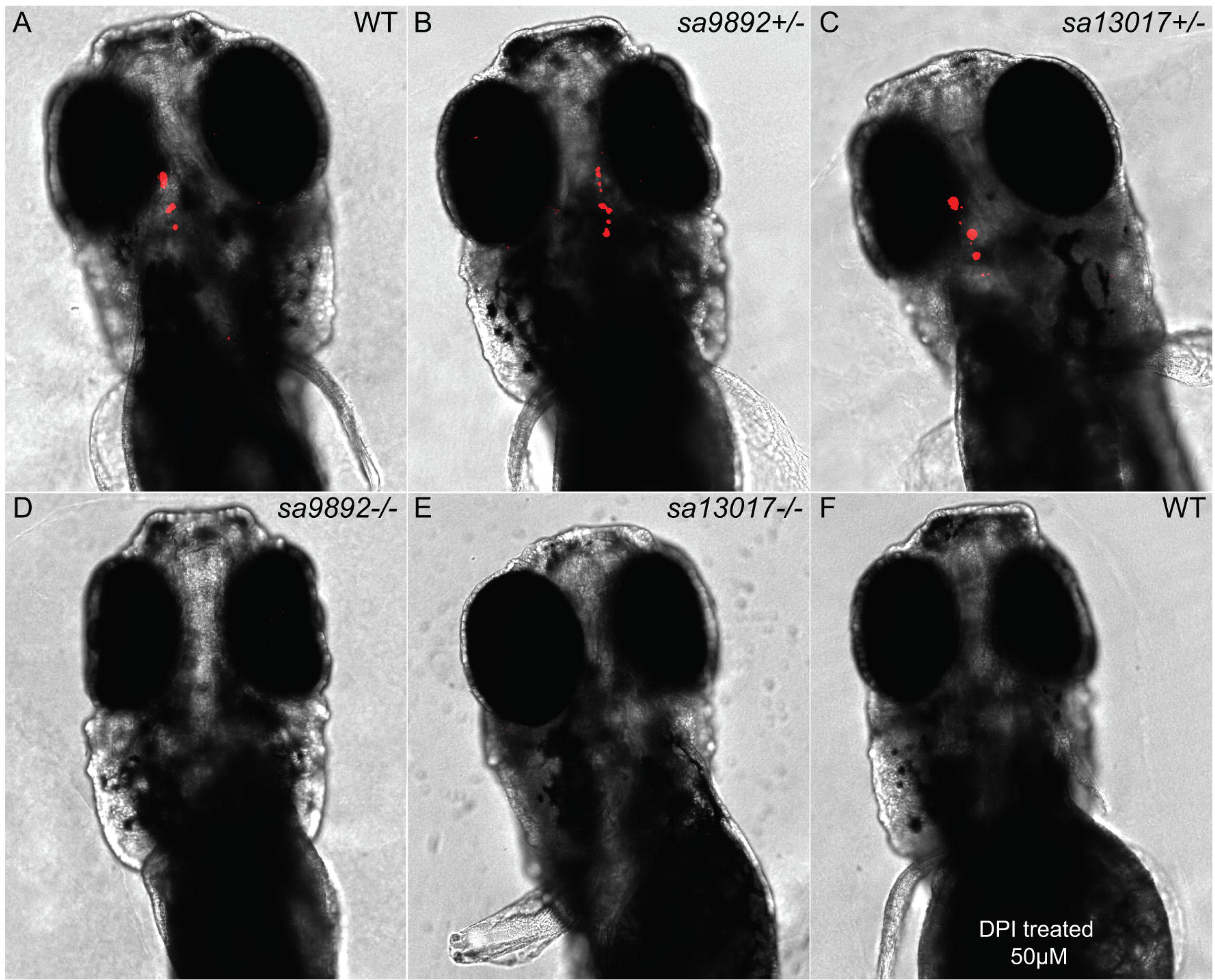
Hypothyroidism is evident among *duox* mutants. At 5dpf, homozygous mutant larvae lack staining bound T_4_ in the thyroid follicles, based on whole-mount fluorescent immunohistochemistry (D–E), which is in sharp contrast to the robust staining observed in wild type and heterozygous siblings (A–C). The NADPH oxidase inhibitor DPI successfully phenocopies *duox* mutations in WT larvae, resulting in an absence of T_4_ detection (F).

### *duox*-Mediated Hypothyroidism Is Responsive To T_4_ Treatment

Among humans, CH responds very well to T_4_ treatment, especially when treatment is initiated as soon as hypothyroidism is suspected (Rahmani et al., 2016). Here we decided to ask whether supplementation of the aquarium water with T_4_ could reverse some or all of the phenotypes observed in the homozygous *duox* mutants. We initiated T_4_ (30nM) treatment of the *duox sa9892^-/-^* and *sa9892^+/-^* animals starting at 11months of age, when all of the phenotypes described previously were already apparent. We found that most of the phenotypes associated with loss of *duox* function could be reversed by treatment with T_4_. Body pigmentation was the first phenotype to be reversed in the treated animals, such that by two weeks after initiation of treatment the *duox sa9892^-/-^* animals became visibly paler than their untreated *duox sa9892^-/-^* siblings (Fig. 7A versus Fig. 7B). The difference in pigmentation was associated with a significant decrease in melanophore density in the homozygous mutant animals treated with T_4_, when compared to the untreated homozygous mutant animals (Fig. 7E-7G). Indeed, the density of melanophores in the treated homozygous mutant animals was similar to that seen in untreated or treated heterozygous mutant animals, suggesting a complete rescue (Fig. 7C-7G). In addition, we found that fin quality improved markedly amongst the treated individuals showing fuller, unbroken fins compared to the ragged fins of the untreated controls (compare Fig. 7A and 7B). Furthermore, after eight weeks of T_4_ treatment we were able to rescue breeding behavior in both sexes. Mutant males and females were able to spawn with wild-type animals or with each other. These mating episodes resulted in the production of 4 clutches of eggs in 4 consecutive weeks. Rescue of fertility was perhaps the most striking outcome of T_4_ treatment.

**Fig. 7.**
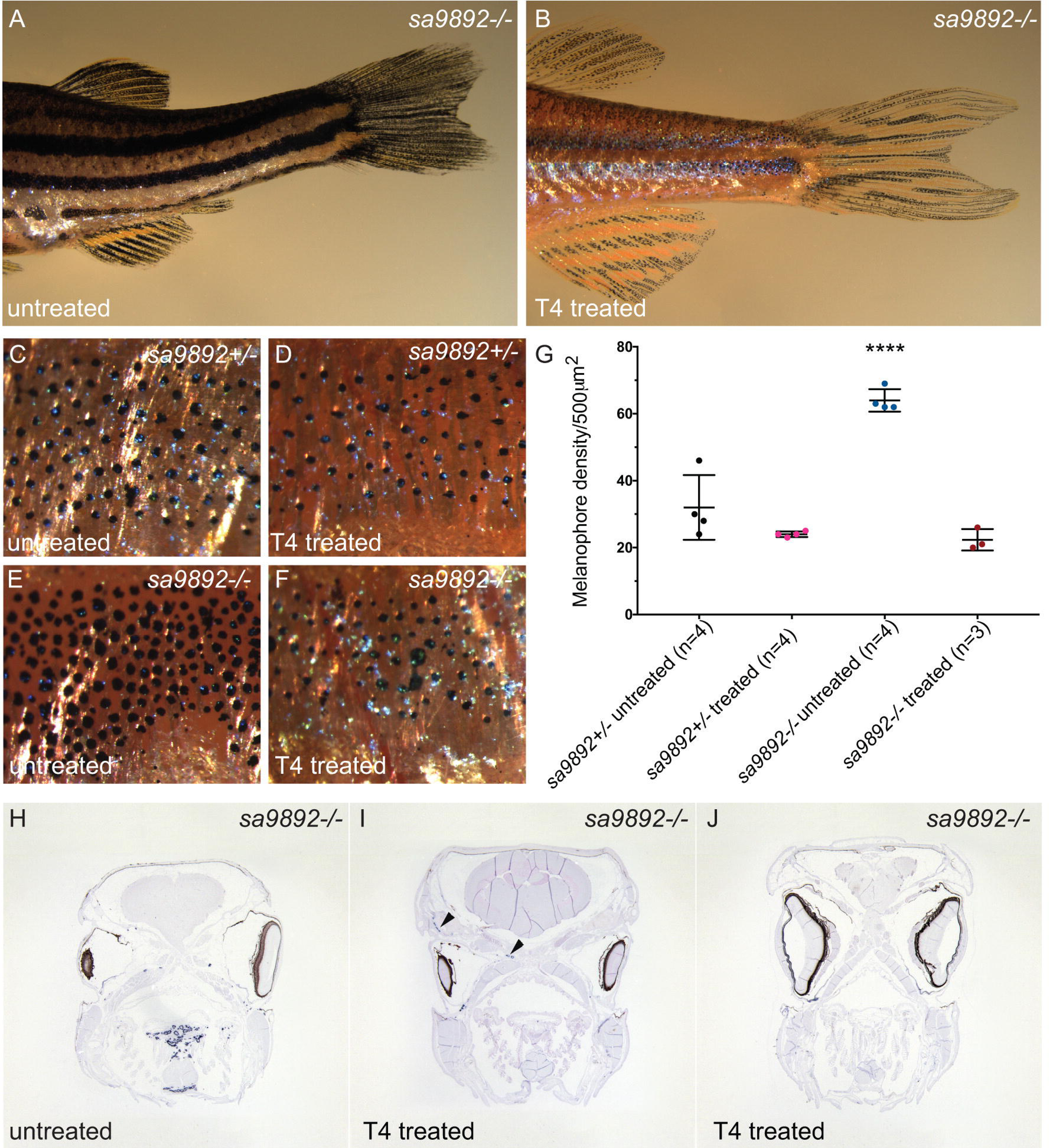
T_4_ treatment alleviates phenotypic anomalies in *duox* mutants. 5X magnification of flank region showing the distribution of melanophores in *sa9892^+/-^* and *sa989^-/-^* siblings (A–D). Pigment changes are evident among T_4_-treated mutants, with a significant reduction in melanophore number (F). Asterisks denote statistically significant differences (Bonferroni’s multiple comparisons test, **** P<0.0001). Treated animals also show an improvement in fin health, compared with untreated controls (G–H). Goitres resolve following T_4_ administration, but small ectopic thyroids are still evident (dark arrows) (I–J).

Having observed a rescue of most of the phenotypes associated with the homozygous *duox* mutants, we wondered whether T_4_ treatment also diminished the size of the thyroid gland in the *duox* mutants. Anecdotally, we had noted that one of the homozygous *duox sa9892^-/-^* mutant animals in the treated group had a small external goitre before treatment, but the goitre resorbed after 2 weeks of treatment. In comparison, a homozygous mutant sibling in the untreated group, that also had an external goitre, showed an increase in the size of the goitre during the course of the experiment (data not shown). This suggested that T_4_ treatment might lead to a diminution in the size of the thyroid glands in the homozygous mutant animals. To confirm whether this was the case, we sectioned and performed ISH for *thyroglobulin* on some of the treated and untreated homozygous mutant animals. We found that treatment led to a dramatic decrease in the thyroid hyperplasia normally associated with the *duox* homozygous mutants, although some of the treated animals did retain some small ectopic *thyroglobulin* staining in the head not seen in wild-type animals, suggesting that these animals had extensive ectopic thyroid follicular tissue prior to treatment (Fig. 7H-7J).

### Methimazole Phenocopies *Duox* Mutant Phenotypes

For final confirmation that the phenotypes found in the homozygous *duox* mutants were due to hypothyroidisms, we asked whether exposure of wild-type fish with the goitrogen methimazole (1mM) phenocopied the homozygyous mutant phenotypes. To counter the influence of already circulating THs, we exposed the adult fish over a three-month period. Treated animals became darker, owing to an increase in the number of melanophores (Fig. 8A-8C). In addition, adult fish treated with methimazole failed to breed, as we had seen in the homozygous duox mutant animals. They also developed external goitres (3 out of 7) (Fig. 8F) and, internally, their thyroid follicles spread dramatically in area (Fig. 8G-8H). This was reminiscent of the observations made in homozygous *duox* mutants (Fig. 8I). Finally, larval WT zebrafish continuously treated with methimazole from between 3hpf and 5hpf onwards showed greatly diminished follicular T_4_ immunostaining, similar to that found in *duox sa9892^-/-^* and *duox sa13017^-/-^* mutant larvae (Fig. 8D-8E).

**Fig. 8.**
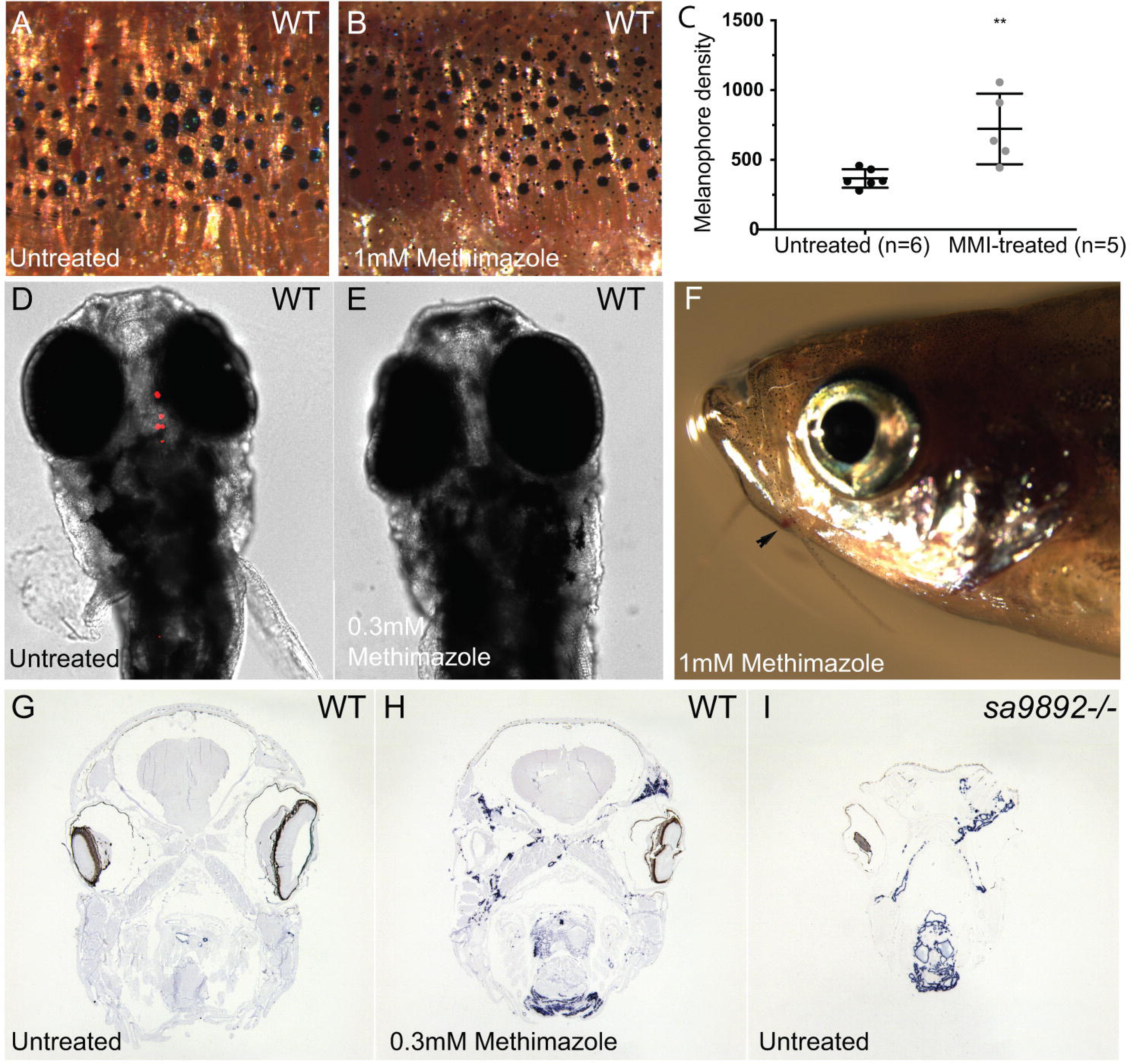
The goitrogen methimazole (MMI) phenocopies *duox* mutations. 5X magnification of flank region showing the distribution of melanophores among MMI-treated and untreated wild type fish. Treated animals have at least two distinct populations of melanophores, based on size (A–B). Pigment change pertaining to melanophore numbers is significant following MMI-treatment (C). (Bonferroni’s multiple comparisons ** P<0.01). MMI leads to loss of bound T_4_ in wild type larvae (D–E) and induces external goitre (F). ISH for thyroglobulin reveals widespread follicular tissue, not limited to the mid-ventral region (H).

## DISCUSSION

The zebrafish has recently emerged as a new, genetically tractable model for investigating the molecular mechanisms underpinning thyroid organogenesis and function (Alt et al., 2006; Elsalini and Rohr, 2003; Guillot et al., 2016; McMenamin et al., 2014; Trubiroha et al., 2018; Wendl et al., 2002). Despite the growing body of work in zebrafish aimed at understanding thyroid development, function and disease, there have been no reports describing the phenotypic consequences of *duox* mutations in adult zebrafish. This is despite the fact that mutations in *DUOX2* and *DUOX1* have been shown to be associated with congenital hypothyroidism in humans for more than a decade (Aycan et al., 2017; Donkó et al., 2014; Jin et al., 2014; Johnson et al., 2007; Kizys et al., 2017; Tonacchera et al., 2009; Vigone et al., 2005). Here, we describe the phenotypes associated with homozygosity of two loss-of-function alleles of zebrafish *duox* in adult fish. Despite the additional round of genome duplication in teleost fish (Taylor et al., 2003), there only exists a single orthologue of *duox* in zebrafish, instead of the two orthologues present in tetrapods (*DUOX1* and *DUOX2*)(Kawahara et al., 2007). Remarkably then, in this instance, zebrafish has less genetic redundancy for this gene than is commonly found in this system. Thus, assessing phenotypes associated with homozygosity of the single *duox* ortholog in zebrafish has allowed us to essentially model the effect of losing function of both *duox* orthologues in tetrapods. This is particularly important as mutations in both *DUOX1* and *DUOX2* in humans have been associated with a more severe form of CH (Aycan et al., 2017), suggesting that *DUOX1*, while normally playing a minor role in TH synthesis in humans, does partially compensate for the loss of *DUOX2* in humans.

Amongst the various phenotypes displayed by the homozygous *duox* mutants, most have been previously observed following pharmacological disruptions in thyroid hormone synthesis or in mutant strains where the hypothalamic–pituitary–thyroid (HPT) axis in zebrafish is affected. For example, goitrogen treatments, thyroid ablation and *tsrh* mutant strains display alterations in pigmentation (McMenamin et al., 2014), similar to those we found in homozygous *duox* mutants. More specifically, thyroid ablated zebrafish have a darker striped pattern due to an increase in the density of melanophores within each stripe (McMenamin et al., 2014), akin to our homozygous *duox* mutants. Another notable phenotype in our homozygous mutants reminiscent of prior findings in goitrogen treated zebrafish was erythema in the proximity of the operculum. Schmidt and Braunbeck (2011) came across a striking histopathological phenotype following treatment of WT zebrafish with the goitrogen, phenylthiouracil (PTU) (Elsalini and Rohr, 2003), wherein treatment resulted in excessive proliferation of blood vessels surrounding the thyroid follicles. This proliferation is attributed to hyperemia resulting from blood aggregation in proximally swollen blood vessels surrounding the thyroid follicles and is concentration-dependent, with the highest concentrations leading to hyperemia (Schmidt and Braunbeck, 2011). Macroscopically, this proliferation of vasculature manifests as erythema, giving a red colour to the entire opercular region. While we did not perform a histological examination of the vasculature, our macroscopic observations are consistent with these reported findings. All our mutants displayed this conspicuous redness of the opercular region. The colouration was most notable amongst younger animals and especially apparent in backgrounds lacking melanophores. Space constraints together with follicular expansion and vascular proliferation in the pharyngeal region could also explain for the flaring opercula observed in some mutants, although this could also be due to thyroid hyperplasia, which was also noted by Schmidt and Braunbeck (2011) in their PTU treated fish.

We were also able to induce this chronic hypothyroid/goitrogenic state in WT animals following treatment with methimazole, resulting in similar phenotypic outcomes. In our reverse experiment, however, it was very interesting to note that while T_4_ treatment of mutants resolved the goitres, some follicular staining remained in ectopic regions. Even so, the overall amount of thyroid tissue was largely diminished. It has been reported that at concentrations ≥25mg/L of PTU, follicular encroachment is found in the gills of zebrafish, suggesting ectopic follicular expansion (Schmidt and Braunbeck, 2011). Follicular expansion following exposure to MMI is attributed to hyperplasia, both, in zebrafish (Schmidt and Braunbeck, 2011) and in frog tadpoles (Hsü et al., 1974). This is regarded to be the first step in compensating for TH production via TSH (Schmidt and Braunbeck, 2011). Concentration dependent increases in the extent of follicular hypertrophy and hyperplasia has also been reported in the fathead minnow (*Pimephales promelas*), when exposed to the thyroid peroxidase inhibitor 2-mercaptobenzothiazole (Nelson et al., 2016). *duox* mutants and MMI-treated WTs presented with amplified *thyroglobulin* expression that showed follicular crowding in the pharyngeal regions and invasion in other ectopic locations. Thyroid dyshormonogeneis (TD) is among the leading causes of CH, and ectopia (ectopic thyroid) is the commonest subtype of TD (De Felice and Di Lauro, 2004). Ectopic thyroid glands have recently been reported in human *DUOX2* mutations wherein scintigraphy revealed submandibular and sublingual thyroid ectopic locations (Kizys et al., 2017).

Fins in teleost fish have garnered interest amongst the scientific community not only due to their capacity for extensive morphological diversity, but also due to their remarkable regenerative capacities (Johnson and Bennett, 1999; Nakatani et al., 2007). Fins are composed of multiple branched and multi-segmented rays covered in a thin layer of epidermal cells. Individual rays consist of a pair of hemirays. Mature hemirays, known as lepidotrichia, are surrounded by a monolayer of osteoblasts that synthesise the bone matrix. With no musculature present, the remainder is made up of mesenchymal cells with nerve fibres and vasculature running along and inside the fin rays. Their remarkable regenerative capabilities underlie not only their ability to regenerate following injury, but also the mechanism by which fins maintain their integrity during normal growth and homeostatic maintenance. Because of the constant growth, renewal and maintenance of the fins, it is relatively uncommon to find animals in aquaria with damaged fins (Wills et al., 2008). Thus, the ragged fins in the *duox* mutants stand out. We have found that the presence of ragged fins is ameliorated, however, by treatment of the mutants with T_4_. Our observations are in line with those in the medaka hypothyroidism mutant, *kmi^-/-^*, which also frequently exhibit damaged or ragged fins (Sekimizu et al., 2007). Furthermore, *kmi^-/-^* animals have also been reported to show delayed regeneration, which can be rescued via exogenous T_4_.

CH has been associated with cephalic and facial defects and developmental neurological abnormalities (Gamborino et al., 2001). Such defects have been attributed to improper development of the cranial neural crest (CNC), which is a transient population of migratory embryonic stem cells. Arising from the neural ectoderm, these cells contribute to a long list of cell types, including bone, cartilage, craniofacial connective tissue, corneal stroma and endothelium, iris stroma, ciliary body stroma and muscles, sclera and the trabecular meshwork of the eye (Barembaum and Bronner-Fraser, 2005; Minoux and Rijli, 2010). An investigation of craniofacial morphogenesis using rats exposed to MMI revealed a 25% reduction in the overall head size throughout gestation (Gamborino et al., 2001). These findings are consistent with observations on craniofacial shape in zebrafish *manet^wp.r23e1^* mutants as well as metronidazole-mediated thyroid ablated transgenics *Tg(tg:nVenus-2a-nfnB)^wp.rt8^*, which have narrower heads than controls (McMenamin et al., 2014). Further evidence on the role of THs has been gathered using pharmacological and morpholino-based approaches in zebrafish larvae. In one study, MMI treatment resulted in reduced head depth and shorter jaw length (Liu and Chan, 2002). In another study, MMI and PTU were found to inhibit pharyngeal arches and ceratohyal cartilage development, while knockdown of *thraa* (thyroid receptor α a) led to malformations in the Meckels and ceratohyal cartilages (Bohnsack and Kahana, 2013). Our observations of the shorter frontal height among *duox sa9892^-/-^* and *sa9892/sa13017* animals are yet another indicator of TH deficiency. The lack of a significant difference in frontal height between the *duox sa13017^-/-^* and the other groups was surprising but may be due to a small sample size. Even among the *duox sa9892^-/-^* animals, while frontal height was significantly different from that of the heterozygotes and WTs, the spread of data itself was quite wide. In contrast, data on *sa9892/sa13017* animals was more clustered and representative of the mean.

*duox* mutants appear to experience several phenotypes associated with retarded growth and development.. These include delayed growth rate, and delayed or incomplete swim bladder morphogenesis and barbel emergence. As development is underway, fish standard length (SL) is subject to both genetic and environmental factors, thus introducing variation amongst siblings. Indeed, environmental influences on SL is clearly apparent as larvae reared in groups show greater variation in SL than larvae raised individually (Parichy et al., 2009). SL is, thus, regarded as a more reliable measure of fish maturation than age (Parichy et al., 2009; Singleman and Holtzman, 2014). Considering the mean values for SL for our groups, it was clearly apparent that at 3 months of age all mutant groups were significantly shorter (i.e. less mature) than their heterozygous or wild type siblings. Remarkably, by 6months, however, the homozygous *duox* mutant fish caught up with the WT and heterozygotes siblings. This suggests that it is not growth per se, but the state of maturation, which is dependent on thyroid hormones. This finding is consistent with findings in non-metamorphosing *Xenopus laevis* tadpoles, which become giants and can live for years in an immature neotenic state (Rot-Nikcevic and Wassersug, 2004). This arrested development, associated with continued growth, has been attributed to a lack of thyroid glands in these animals (Rot-Nikcevic and Wassersug, 2004). In fish, definitive evidence of thyroid hormone insufficiency causing metamorphic stasis is well appreciated from studies on flatfish. Larvae of the summer flounder (*Paralichthys dentatus*), when treated with thiourea, do not develop beyond early metamorphic climax (Schreiber and Specker, 1998). Likewise, olive flounder (*Paralichthys olivaceus*) larvae treated with the goitrogen, thiourea, enter metamorphic stasis and become giant larvae (Inui and Miwa, 1985). Although metamorphosis among the roundfish is less dramatic, several examples illustrate the dependence of metamorphosis on THs. Thiourea treatment was found to arrest metamorphosis in the coral trout grouper (*Plectropomus leopardus*) (Trijuno et al., 2002) orange-spotted grouper (*Epinephelus coioides*) (de Jesus et al., 1998) and the red sea bream, (*Pagrus major*) (Hirata et al., 1989). Meanwhile, the pesticide chlorpyrifos, reported to cause reductions in serum concentrations of T_4_ and T_3_ (Slotkin et al., 2013), was recently found to prevent metamorphic completion in the convict surgeonfish (*Acanthurus triostegus*) (Holzer et al., 2017). In zebrafish, a 1mM concentration of methimazole inhibited the larval to juvenile transition (Brown, 1997). However, larvae treated with a concentration of 0.3mM eventually escaped the inhibition and continued development. While our *duox* mutants eventually reach normal adult size, this might be associated with an incomplete metamorphic or immature state. Alternatively, there may be some genetic redundancy present in zebrafish, whereby a different source of H_2_O_2_ in the thyroid follicles might be capable of partially compensating for the loss of *duox* function. Indeed, another NOX isoform, NOX4, has been described in human thyrocytes. Unlike Duox though, NOX4 generates H_2_O_2_ in the intracellular compartment (Weyemi et al., 2010). It may thus be important to generate double mutants for *duox* and *nox4* to determine the contribution of Nox4 in thyroid hormonogenesis.

In the zebrafish, swim bladder inflation is dependent on THs (Godfrey et al., 2017; Liu and Chan, 2002). The posterior chamber of the swim bladder inflates around 4.5dpf while the anterior chamber inflates by 21dpf (Winata et al., 2009). These events appear to coincide with peaks in whole body T_3_ at 5dpf and 10dpf and T_4_ at 21dpf (Chang et al., 2012). Previously, it was found that swim bladder inflation was significantly delayed in thyroid-ablated zebrafish, where the anterior chamber of the bladder inflated ~50dpf, compared to ~20 days in controls (McMenamin et al., 2014). Similar observations were also made in thyroid-ablated *Danio albolineatus* (McMenamin et al., 2014). There also exists sufficient evidence of how pharmacologically disrupted thyroid processes affect swim bladder inflation. Ecological assessments of aquatic pollutants often employ key morphological events during fish development as predictive approaches. 2-Mercaptobenzothiazole (MBT), commonly used for rubber vulcanization, is found to occur in environmental water bodies. MBT is a potent TPO inhibitor and its role was recently examined in swim bladder inflation in the fathead minnow (*Pimephales promelas*) (Nelson et al., 2016) and zebrafish (Stinckens et al., 2016). Among minnows, larvae continuously exposed to MBT showed a concentration-dependent decrease in anterior lobe size (Nelson et al., 2016). Meanwhile, MBT-treated zebrafish larvae were reported to fare worse than minnows, where 22% larvae exposed to the highest concentration failed to inflate the anterior chamber (Stinckens et al., 2016). Interestingly, even though both species belong to the order, Cypriniformes, a compensatory T_4_ response has been reported in the fathead minnow at 21dpf (Nelson et al., 2016) but not in the zebrafish (Stinckens et al., 2016), suggesting species-specific differences. Our homozygous *duox* mutant animals also displayed a significant delay in anterior chamber inflation, suggesting that Duox is essential for this process, likely through its role in thyroid hormone synthesis.

Barbels are yet another easily observable phenotypic trait influenced by thyroid hormones. In zebrafish, both pairs develop as epithelial buds around 30-40dpf, following the emergence of pelvic fin rays (Hansen et al., 2002; Parichy et al., 2009). Thus far, only one study has reported barbel emergence to be influenced by THs. Thyroid ablation via Mtz of *Tg(tg:nVenus-2a-nfnB)* of *D. rerio* and *D. albolineatus* resulted in the absence of sensory barbels (McMenamin et al., 2014). Similarly, *manet* mutants also lacked barbels (McMenamin et al., 2014). Our homozygous *duox* mutants also show impairment of barbel emergence, consistent with their hypothyroid state. However, it is notable that a subset of the *sa9892-/-* mutants eventually did grow barbels, similar to their body length catch-up phenotype.

In humans, thyroid dysfunction during pregnancy has been positively associated with adverse maternal/foetal outcomes, including infertility, miscarriage, pre-eclampsia, pre-term (before 37weeks) birth and maternal thyroid dysfunction postpartum (Hernández et al., 2018; Stagnaro-Green et al., 2011; Velasco and Taylor, 2018). TH is essential for early development and maturation of the foetal brain and maternal transfer of TH is especially important during the first trimester (Cooper and Biondi, 2012) since the embryo does not begin synthesizing THs until 12-13 weeks into gestation (Casey and Leveno, 2006). The British Thyroid Foundation suggests prescribing levothyroxine to hypothyroid women trying to conceive in order to address these negative consequences of hypothyroidism on fertility and pregnancy. Intriguingly, we also noted significant defects in fertility in both sexes in our homozygous *duox* mutants. Although we do not currently know the reason for infertility in the *duox* mutants, a potential cause may be due to failure in mating behavior as a consequence to our observed effects on pigmentation in the mutants. It has previously been noted that, in zebrafish, both sexes experience changes in their stripes and interstripe colours in the morning, a process termed ephemeral sexual dichromatism, during mating and spawning (Hutter et al., 2012). Another study reported that females utilize yellow colouration for sex recognition (Hutter et al., 2011). This ties in well with xanthophore deficiency reported in thyroid ablated, hypothyroid zebrafish (McMenamin et al., 2014), and by extension, the *duox* mutants. However, it is notable that casper strains of zebrafish, which lack xanthophores altogether, can successfully breed (White et al., 2008). Thus, there may be additional factors that may be contributing to infertility in *duox* mutants.

Associations between thyroid status and reproduction in teleosts have been previously reviewed (Cyr and Eales, 1996). Four physiological pre-requisites have been recognized as essential to spawning behavior and fertility in fish: 1) the completion of vitellogenesis in the ovaries, 2) maturation and ovulation of oocytes stimulated by pituitary luteinizing hormone (LH), 3) completion of spermatogenesis, and 4) sufficient production and storage of milt (seminal plasma and mature sperm) in the sperm duct. These are largely regulated by the endocrine system (Kobayashi et al., 2002). T_3_ enhances the response of the ovarian follicles to gonadotropins, thus facilitating secretion of 17β estradiol (Cyr and Eales, 1988). This regulates the production of vitellogenin by the liver, and in studies on Great Lakes salmonids it has been suggested that lowered T_3_ levels may impair oocyte production (Leatherland and Barrett, 1993). In the fathead minnow (*Pimephales promelas*), methimazole treatment led to a reduction of the cortical alveolus oocytes, relative to control females. Meanwhile, in post-spawning males, control animals showed an increase in the number of spermatozoa and a decrease in the number of spermatogonia. This increase in spermatozoa was not observed in methimazole-treated cohorts, suggesting that hypothyroidism affects spermatogenesis (Lema et al., 2009). Among the African sharptooth catfish (*Clarias gariepinus*), pre-spawining males treated with thiourea were shown to have narrower seminiferous tubules and fewer spermatozoa (Swapna et al., 2006). Intriguingly, hypothyroidism in humans has also been associated with impaired spermatogenesis and sperm abnormalities (La Vignera and Vita, 2018). We have found that fertility in our homozygous *duox* mutants can be restored in both sexes and we can successfully raise offspring to adulthood from a cross between a mutant male and WT female. This is in line with previous observations on growth-retarded (*grt*) mice. *grt* mice have autosomal recessive hypothyroidism, with females suffering lifelong infertility and males gradually acquiring fertility. When treated with THs, *grt* females showed an increase in the size of their uteri and ovaries, which was comparable with heterozygous and WTs. Furthermore, they engaged in copulatory behavior and were able to conceive and deliver pups (Hosoda et al., 2008). Zebrafish *duox* mutants thus provide an excellent model to investigate the consequences of human CH associated with mutations in *DUOX1* and *DUOX2*, and the mechanisms by which treatment with THs, even in adults, can restore many of the defects caused by chronic hypothyroidism, including restoration of fertility in both males and females.

## CONCLUSIONS

Overall, we found that homozygous mutants display a number of phenotypes, which can be ascribed to hypothyroidism, including growth retardation, pigmentation defects, ragged fins, thyroid hyperplasia and external goitre. By and large, the growth retardation defect is not permanent, as fish continue to grow despite being chronically hypothyroid, and ultimately catch up with their euthyroid heterozygous and wild type siblings. This contrast with findings in humans suffering from hypothyroidism, which remain growth retarded unless T_4_ treatment is initiated within weeks after birth. Most other phenotypes associated with chronic hypothyroidism in the *duox* mutant fish were rescued by T_4_ treatment, even if supplementation was not initiated until adulthood. These include recovery of fertility, return to normal pigmentation, improvement in fin morphology and return to normal size thyroid glands. In summary, *duox* mutant zebrafish provide a new and potentially powerful system to understand the consequences of chronic congenital hypothyroidism on growth and maintenance of body physiology, as well as the mechanisms of recovery of normal physiology following thyroid hormone supplementation. Thus, our *duox* mutant fish appear to be in a chronic hypothyroid / goitrogenic state, as indicated by their external goitres as well as internal expansion of *thyroglobulin* expressing tissue.

## MATERIAL AND METHODS

### Ethics Statement

All experiments involving animals were approved by the local ethics committee and the Home Office.

### Animals and Husbandry

Adults and larvae were used in this study. The zebrafish (*Danio rerio*) wild-type (WT) line used was AB. Mutant lines used were *duox sa9892* and *duox sa13017* (Kettleborough et al., 2013) and were obtained from the European Zebrafish Resource Center (EZRC). Compound heterozygotes for these mutant alleles were generated in-house. Both *duox* alleles were also crossed into *nacre* (*nac^w2^*) (Lister et al., 1999) and *casper* (White et al., 2008) strains for visualising larval thyroid follicles, swim bladder and adult erythema. In all cases, embryos were raised in sea salts (Sigma-Aldrich S9883) medium containing 0.0001% methylene blue until 5 days post-fertilisation (dpf) and then transferred to the system where they were maintained at a temperature of 28°C, pH 7.4, constant salinity and a 14:10 photoperiod.

### PCR and Genotyping

Genomic DNA was extracted from caudal fin clips or whole larvae using lysis buffer, in a thermal cycler. The conditions for this procedure were 2hrs at 55°C, 10mins at 95°C and a hold (if necessary) at 12°C. PCR was performed using ExTaq DNA polymerase (TaKaRa RR001A) with the following primer pairs: for the *duox sa9892* allele, forward 5’-ACGAGGTACACAACTCAAGCTG-3’ and reverse 5’-GACGTTCAAAGCGAAACCTGAC-3’; for the *duox sa13017* allele, forward 5’-TGGTACACCATTTGAGGATGTGA-3’ and reverse 5’-ACACCCACCATAGAGGTCTCT-3’. PCR conditions were as follows: 36 cycles at 94°C for 30s, 55°C for 30s and 72°C for 30s. Samples were subject to Sanger sequencing (GATC Biotech). Sequencing primers used were 5’-CTTGGTCTGCCTTTGACGAAGT-3’ for the *duox sa9892* allele and 5’-GTGACTCAAGTCAGAACAGGTC-3’ for the *duox sa13017* allele. Siblings were stage-matched, phenotypically WT, heterozygous and homozygous animals obtained by crossing heterozygous carriers.

### Whole mount immunofluorescence

Zebrafish larvae, at 5dpf, were fixed overnight in 4% phosphate-buffered paraformaldehyde (PFA) (Sigma Aldrich), at 4°C. This was followed by 15mins of dehydration in 100% methanol. Larvae were then transferred to fresh 100% methanol and stored at -20°C until usage. Larvae were gradually rehydrated to PBST, treated with 10μg/ml proteinase K (Roche) for 30min, briefly rinsed in PBST, and postfixed in 4% PFA for 20min. Following further rinsing in PBST, larvae were immersed in blocking buffer (PBST containing 1% dimethylsulfoxide, 1%BSA, 5% horse serum and 0.8% Triton X-100) for 2h. This was followed by overnight incubation, at 4°C, in blocking buffer containing the primary antibody (1:1000) against thyroxine (T_4_) (Biorbyt orb11479). Overnight incubation was followed by several wash steps in PBST containing 1% BSA. Larvae were then incubated overnight, at 4°C, in blocking buffer containing the secondary antibody (1:250) Alexa Fluor 568 (Invitrogen A11057) (Opitz et al., 2011). Stained larvae were washed in PBST, imaged and then subject to genotyping.

### Histology and *in situ* hybridization

In situ hybridization on sections of adult zebrafish was performed essentially as described (Paul et al., 2016). Briefly, adult zebrafish were fixed whole in 4% PFA for a week followed by a decalcification step in 20% EDTA, for 10 days. Animals were then cut at the operculum and mid trunk level and processed in a Leica TP1050 tissue processor in preparation for paraffin embedding. The embedding station used was a Leica EG1150H. The cut face of the tissue was oriented towards the leading edge of the paraffin block and sectioned at 5μm thicknesses on a LeicaRM2255 microtome. Sections were arranged and held on Superfrost Plus™ slides (Thermo Fisher Scientific). Alternating sections were then taken forward for Haematoxylin and Eosin (H&E) staining and *in situ* hybridization (ISH). Sections were put through H&E staining via a Varistain 24-4 carousel (Thermo Shandon). *Thyroglobulin* (*tg*) cDNA used for riboprobe synthesis was amplified using forward 5’-AGGTGGAGAATGTTGGTGTG-3’ and reverse 5’-CTCCAACTCTGGCAATGACT-3’ primers. Digoxigenin-labelled probes were synthesised *in-vitro* using a MEGAscript ^®^T7 kit (Ambion).

### Body length and melanophore counts

For measuring body length, all fish were briefly anaesthetised in 0.02% MS 222 (tricaine) (Sigma-Aldrich). They were then transferred onto an agarose bed in a petridish and imaged at 0.73X magnification. For each fish two or three images were captured in order to include the entire length including the caudal fin. These part images were stitched together in Adobe^®^ Photoshop to obtain a single image. The ‘ruler’ tool and ‘analyse measurement’ command in Photoshop CS5 were used on these images to calculate the length from the tip of the mandible to the caudal peduncle.

For determining melanophore density, all fish were treated with epinephrine to contract pigment granules. To obtain a 1mg/ml working solution, 0.1g of epinephrine was dissolved into 100ml of a 0.01% tricaine solution. Epinephrine is only partially soluble in water and thus, the solution was filtered to obtain a clear filtrate. The solution turns pink-orange during filtration. Fish were treated for 5 minutes in this working solution. They were then transferred onto an agarose bed in a petridish and imaged at three locations on the lowermost continuous stripe extending from the operculum to the tail tip. A 2X or 5X magnification was used. The ‘multi-point’ tool on FIJI was used to manually count melanophores in a 2mm^2^ area for each image. All images were acquired on a Leica MZ16FA fluorescence stereomicroscope with a DC490 camera.

### Pharmacological treatments

For rescuing mutant phenotypes, a 12-week treatment with T_4_ (Sigma-Aldrich) was sustained in a closed system that closely resembled aquarium conditions. 4 groups-*sa9892^+/-^* untreated, *sa9892^+/-^* treated, *sa9892^-/-^* untreated and *sa9892^-/-^* treated-were subject to this regime, with each group comprised of 4 adult fish. T_4_ was added tri-weekly, at a concentration of 30nM. Water was changed three times each week.

For phenocopy experiments, a 12-week treatment with methimazole (Sigma-Aldrich) was administered, once again, in a closed system simulating aquarium conditions. This regime was applied to 6 WT adult fish, while 6 untreated WT animals comprised the control group. Methimazole was added tri-weekly, at a concentration of 1mM. Water was changed three times each week.

For immunostaining, WT larvae were treated with methimazole (0.3mM) from 0dpf. Animals were then fixed at 5dpf and stained for T_4_ as described above.

### Statistical analyses

GraphPad Prism 7 was used for statistical testing. Column statistics and analyses of variance were implemented for all data sets. For column statistics, we calculated median, SD, SEM, confidence intervals and Gaussian distribution. The D’Agostino and Pearson test was used to check for Gaussian distribution. One-way ordinary ANOVA was used to analyse variance. Differences were considered significant at P<0.0001. Bonferroni’s multiple comparisons test was used to compare means between groups.

## Acknowledgements

We would like to thank Peter Walker, Grace Bako and Natalie Partington, at the Core Histology Facility, for their help with histological sectioning and staining. We would also like to thank the aquarium staff in the BSF unit for their care and support of the fish. We also extend our gratitude to Simone Schindler, University of Exeter, for her guidance on performing ISH on tissue sections. Finally, we thank Sabine Cotagliola and Pierre Gillotay, Université Libre de Bruxelles, Belgium, for their advise with the T_4_ antibody protocols in larvae.

## Competing Interests

The authors declare no competing interests.

## Author Contributions

Conceptualization: E.A., K.C., S.I.; Methodology: K.C., S.I.; Validation: K.C.; Formal analysis: K.C.; Investigation: K.C., S.I.; Writing - original draft: K.C.; Writing - review & editing: E.A.; Visualization: E.A.; Supervision: E.A..; Project administration: E.A.; Funding acquisition: E.A.

## Funding

This work was supported by a PhD studentship from The Scar Free Foundation (K.C.) and an MRC Research Project Grant (MR/L007525/1) (S.I., E.A.).

